# Experimental evolution of *Pseudomonas aeruginosa* to colistin in spatially confined microdroplets identifies evolutionary trajectories consistent with adaptation in microaerobic lung environments

**DOI:** 10.1101/2023.06.12.544597

**Authors:** Saoirse Disney-McKeethen, Seokju Seo, Heer Mehta, Karukriti Ghosh, Yousif Shamoo

## Abstract

Antibiotic resistance is a continuing global health crisis. Identifying the evolutionary trajectories leading to increased antimicrobial resistance can be critical to the discovery of biomarkers for clinical diagnostics and new targets for drug discovery. While the combination of patient data and *in vitro* experimental evolution has been remarkably successful in extending our understanding of antimicrobial resistance, it can be difficult for *in vitro* methods to recapitulate the spatial structure and consequent microenvironments that characterize *in vivo* infection. Notably, in cystic fibrosis (CF) patients, changes to either the PmrA/PmrB or PhoP/PhoQ two-component systems have been identified as critical drivers for high levels of colistin and polymyxin resistance. When using microfluidic emulsions to provide spatially structured, low-competition environments, we found that adaptive mutations to *phoQ* were more successful than *pmrB* in increasing colistin resistance. Conversely, mutations to *pmrB* were readily identified using well-mixed unstructured cultures. We found that oxygen concentration gradients within the microdroplet emulsions favored adaptive changes to the PhoP/PhoQ pathway consistent with microaerobic conditions that can be found in the lungs of CF patients. We also observed mutations linked to hallmark adaptations to the CF lung environment, such as loss of motility (*fleQ, fliC, fleS, flg, flh*, and *fleQ*) and loss of O antigen biosynthesis (*wbpL*). Mutation to *wbpL*, in addition to causing loss of O antigen, was additionally shown to confer moderately increased colistin resistance. Taken together, our data suggest that distinct evolutionary trajectories to colistin resistance may be shaped by the microaerobic partitioning and spatial separation imposed within the CF lung.

**Importance:** Antibiotic resistance remains one of the great challenges confronting public health in the world today. Individuals with compromised immune systems or underlying health conditions are often at an increased for bacterial infections. Patients with Cystic Fibrosis (CF) produce thick mucus that clogs airways and provides a very favorable environment for infection by bacteria that further decrease lung function and, ultimately, mortality. CF patients are often infected by bacteria such as *Pseudomonas aeruginosa* early in life and experience a series of chronic infections that, over time, become increasingly difficult to treat due to increased antibiotic resistance. Colistin is a major antibiotic used to treat CF patients. Clinical and laboratory studies have identified PmrA/PmrB and PhoP/PhoQ as responsible for increased resistance to colistin. Both have been identified in CF patient lungs, but why, in some cases, is it one and not the other? In this study, we show that distinct evolutionary trajectories to colistin resistance may be favored by the microaerobic partitioning found within the damaged CF lung.

## Introduction

The rise of antibiotic resistance is one of the most pressing threats to global health in the 21st century. *Pseudomonas aeruginosa* is a pathogen of particular concern due to the severe infections it causes in individuals with cystic fibrosis (CF) and immunocompromised patients, its intrinsic resistance to a number of widely used antibiotics, and the ease with which it acquires resistance (1, 2). Clinical resistance to colistin (polymyxin E), often the last resort antibiotic used to treat infections, is becoming increasingly common and highlights the importance of understanding how *P. aeruginosa* acquires colistin resistance and what factors influence specific evolutionary trajectories to resistance (3). In addition to increased antibiotic resistance, there are a number of hallmark phenotypic adaptations to the CF lung environment, including increased biofilm production, loss of motility, and loss of O antigen biosynthesis (4–7). Another defining characteristic of chronic lung infection is the genetic and phenotypic diversity of bacterial populations (8, 9). The CF lung is a spatially structured, heterogeneous environment where limited bacterial intermixing increases the number and type of evolutionary trajectories available to bacterial populations (8, 10). Regionally isolated areas of the lung also vary in nutrients, oxygen, inflammation damage, and antibiotic concentration during treatment which further increases the number of available niches (10).

Experimental evolution is a powerful tool for analyzing the molecular basis of how populations evolve (11, 12), and multiple studies have used experimental evolution to study how *P. aeruginosa* acquires colistin resistance (13–15). Perhaps the most well-established resistance pathways are through the *pmrAB* and *phoPQ* Two-Component Systems (TCS), which have been found to confer resistance in both experimentally evolved strains and clinical isolates (13–17). However, experimental evolution to antibiotics is typically done using serial flask transfer or bioreactors, both of which are well-mixed environments that lack the spatial structure necessary for diverse populations. In these bulk culture environments, bacteria are competing for a shared pool of resources which favors fast growth. Slower-growing resistant strains that might be viable *in vivo* are outcompeted, leading to a loss of diversity. Performing experimental evolution under spatially segregated conditions has a significant impact on the evolutionary outcomes (18), with previous studies utilizing biofilm transfer methods finding populations exhibited significant diversification as well as adaptations similar to those observed in isolates from CF patients (19–21). Previously, we have shown that spatially segregated microdroplet emulsions can identify evolutionary trajectories that were not seen in bulk cultures (22).

In this study, *P. aeruginosa* strain PAO1 was evolved to colistin resistance in microdroplet emulsions to study how resistance is acquired in spatially structured environments and what environmental factors might influence the success or failure of specific evolutionary trajectories. We hypothesized that in environments with spatial segregation, we would identify mutations involved in either colistin resistance, environmental adaptation, or both that were not observed in unstructured batch cultures such as a well-mixed flask. Indeed, we noted significantly different evolutionary trajectories in well-mixed batch cultures versus those of the microdroplet emulsions. In microdroplet environments, changes to the *phoPQ* TCS were strongly favored over changes in the *pmrAB* TCS. While changes to both TCS’s have been identified in CF patients, it is unclear whether mutations in either *phoPQ* or *pmrAB* are more advantageous in different contexts. Our data show that adaptive changes in *pmrB* have a significantly higher fitness cost in the microaerobic conditions that might be found in certain regions of the CF lung. From experimental evolution in microemulsions that had the highest levels of spatial confinement, we also identified changes characteristic of chronic, late-stage CF lung disease (23), namely mutations in genes previously linked to a loss of motility (*fleQ, fliC*) and mutations linked to loss of O antigen biosynthesis (*wbpL*). Strikingly, the loss-of-function mutation in *wpbL* also led to a modest increase in colistin resistance that was successful as an adaptive change early in the progression toward very high colistin resistance.

## Results

### Evolutionary timelines to colistin resistance identified by experimental evolution differ as a result of environmental/spatial structure

To study how colistin resistance is acquired in spatially segregated environments, we evolved populations of *P. aeruginosa* PAO1 (Ancestor strain) to colistin at multiple levels of spatial segregation and in two media conditions. The different populations and growth conditions are summarized in **Figure 1**. To investigate the role of spatial structure in shaping the early stages of adaptation to colistin, we confined cells in microdroplets at different starting cell densities. Populations denoted as λ0.5 had an average of 0.5 cells/microdroplet in 46 ± μm diameter microdroplets, whereas λ91 had an average of 91 cells/microdroplet in 143 ± 5.6 μm diameter microdroplets. The average number of bacteria per microdroplet (λ) was controlled by droplet size and culture dilution factor (24). The λ0.5 populations have far less competition than the λ91 or well-mixed flasks. The λ91 populations were used to provide a more batch-like set of populations in microdroplets that could be compared to the more widely used batch conditions of earlier studies (13–15). λ91 microdroplet populations for experimental evolution were also performed in rich media (LB-BHI) and the lung-sputum simulant SCFM (25). In cases where expansion proceeded slowly due to increases in colistin concentration, the colistin gradient was maintained at the same level until the populations expanded sufficiently to restore the mutation supply, and allow for genomic DNA isolation before progression to the next concentration of colistin. Adaptation to 3–3.5 μg/ml colistin in the λ91 and flask-transfer formats were comparable (6-7 passages), while the λ0.5 populations required 22-25 passages (**S1**). The much longer adaptation required for λ0.5 populations is the result of the much smaller carrying capacity of the 50 μm diameter microdroplets (∼200 cells vs ∼2000 in 143μm microdroplets) that decreases the expansion rate of adaptive lineages (**S2**).

**Figure 1.**
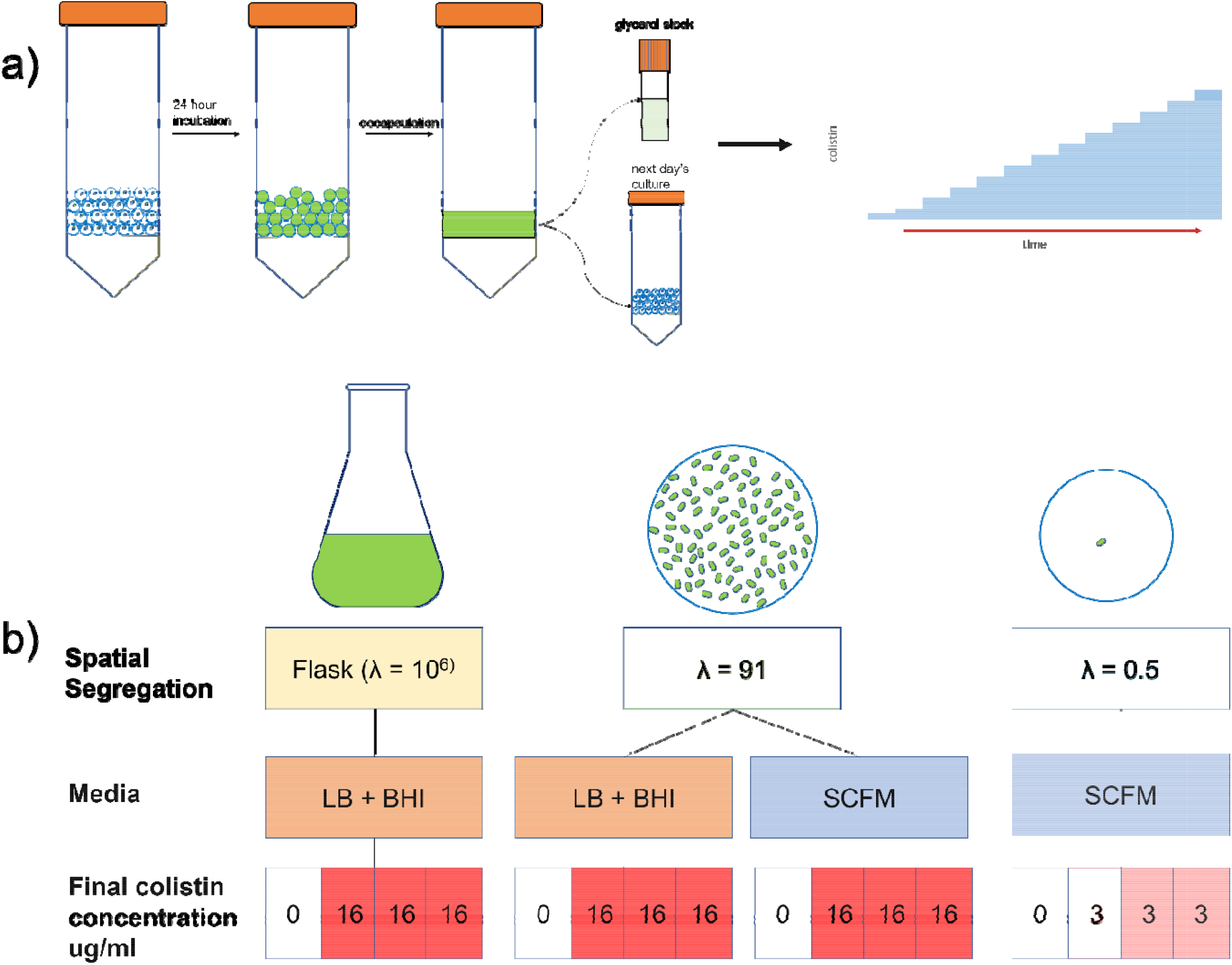
Design of experimental evolution environments. a) experimental evolution in microdroplets that provide controlled spatial structure (green represents growth of cells and yellow is emulsion oil layer produced during encapsulation) λ is the average number of cells in each spatial structure at t=0. b) overall schema for experimental evolution shows the different evolution conditions for each population

### Longitudinal whole genome sequencing of microdroplet populations during the early stages of adaptation shows that *pmrAB* mutations occur early in flask populations but do not emerge in microdroplet populations

Longitudinal whole genome sequencing was performed for all populations to investigate how allele frequencies changed over time as colistin concentration was increased. We investigated the initial phases of colistin adaptation across flask, λ91, and λ0.5 conditions by comparative genomic analysis for all populations up to 3 to 3.5 μg/ml colistin, which is above the minimum inhibitory concentration (MIC) of PAO1 for colistin (1-2 μg/ml) (14). **Fig 2** shows the mutations present in all populations at 3-3.5 μg/ml colistin.

**Fig 2.**
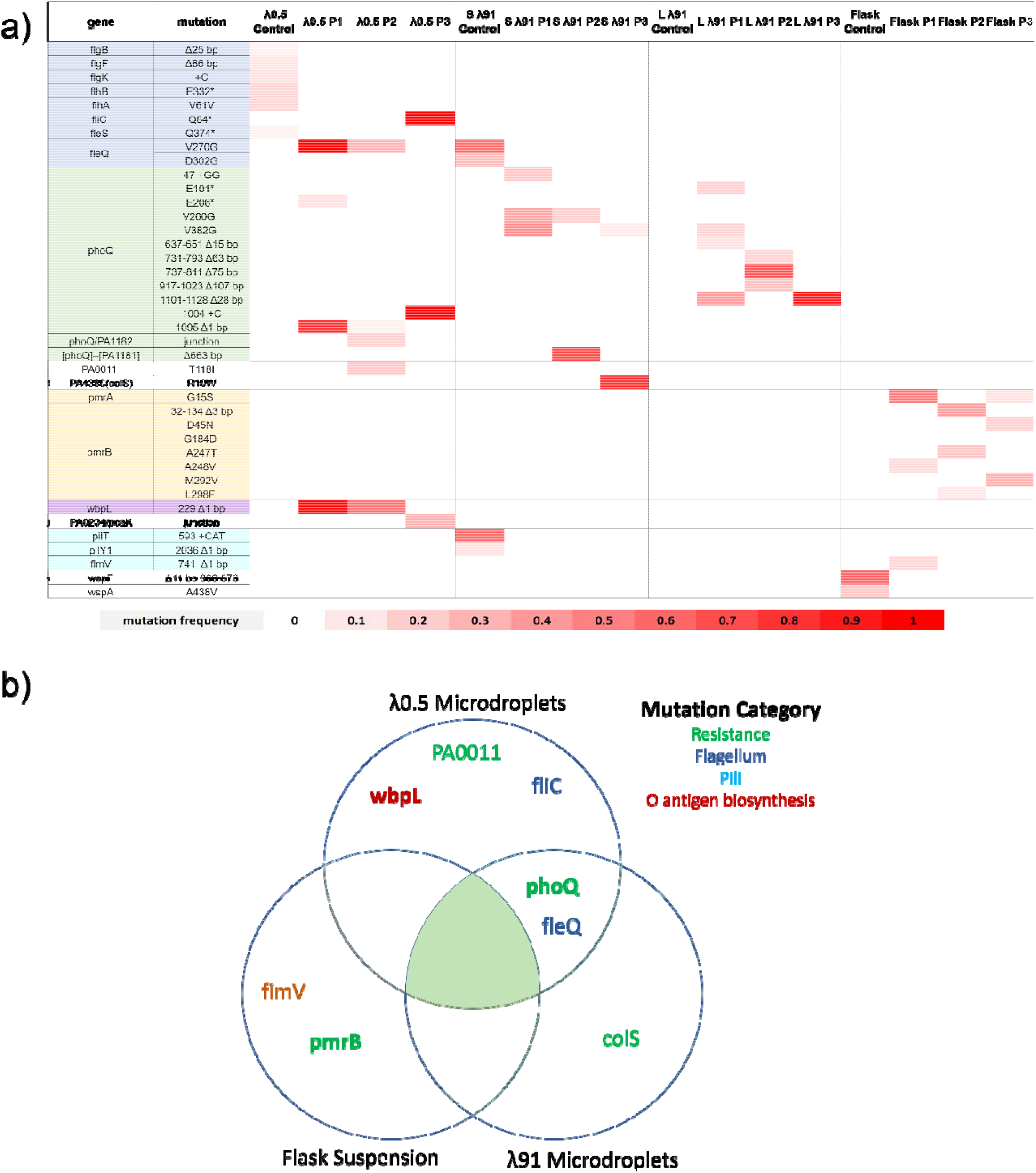
Experimental evolution identified distinct adaptive alleles for the microdroplet versus flask environments. a) heatmap of mutations present in all populations at 3μg/ml colistin shows distinct patterns of mutations in the final populations for the different conditions. For example, changes to *pmrA* or *pmrB* were not identified in any of the microdroplet populations. b) Venn diagram showing significant genes that were mutated above 15% in endpoint populations across different conditions and color coded based on known functions. Mutations associated with O antigen biosynthesis and flagellar motility were also present only in microdroplet conditions.

In flasks, either *pmrB* or *pmrA* mutations were the first to appear on Days 4-5 at around 1-2 μg/ml colistin (**Fig 3**.). Populations 2 and 3 had multiple pmrB mutations, while Population 1 had a dominant mutation (53% (G15S)) in *pmrA* and a lower frequency mutation in *pmrB* (15% (A248V)) by Day 6. In contrast, none of the λ91 populations had mutations in *pmrB*, but all had mutations in *phoQ*, most often deletions, such as the Δ28 bp deletion of base pairs 101□1128 in Populations 1 and 3 of the λ91 LB+BHI condition. (**Fig 2a)**. In Population λ91 SCFM 3, there was an additional mutation in *colS*, another two-component sensor implicated in colistin resistance (26). In the λ0.5 experimental populations, there were mutations in *phoQ, wbpL, fleQ*, and *fliC*, but again not in *pmrA* or *pmrB* (**Fig 2**). This result was unexpected as phoPQ and pmrAB are generally thought to similarly mediate colistin resistance as they both confer resistance through regulation of the *arn* operon (27). Mutations in both the *pmrAB* and *phoPQ* TCS are known to confer colistin resistance in both laboratory evolution and clinical settings (16, 17), although previous continuous culture laboratory experiments mainly identified *pmrAB* mutations (13, 15).

**Figure 3.**
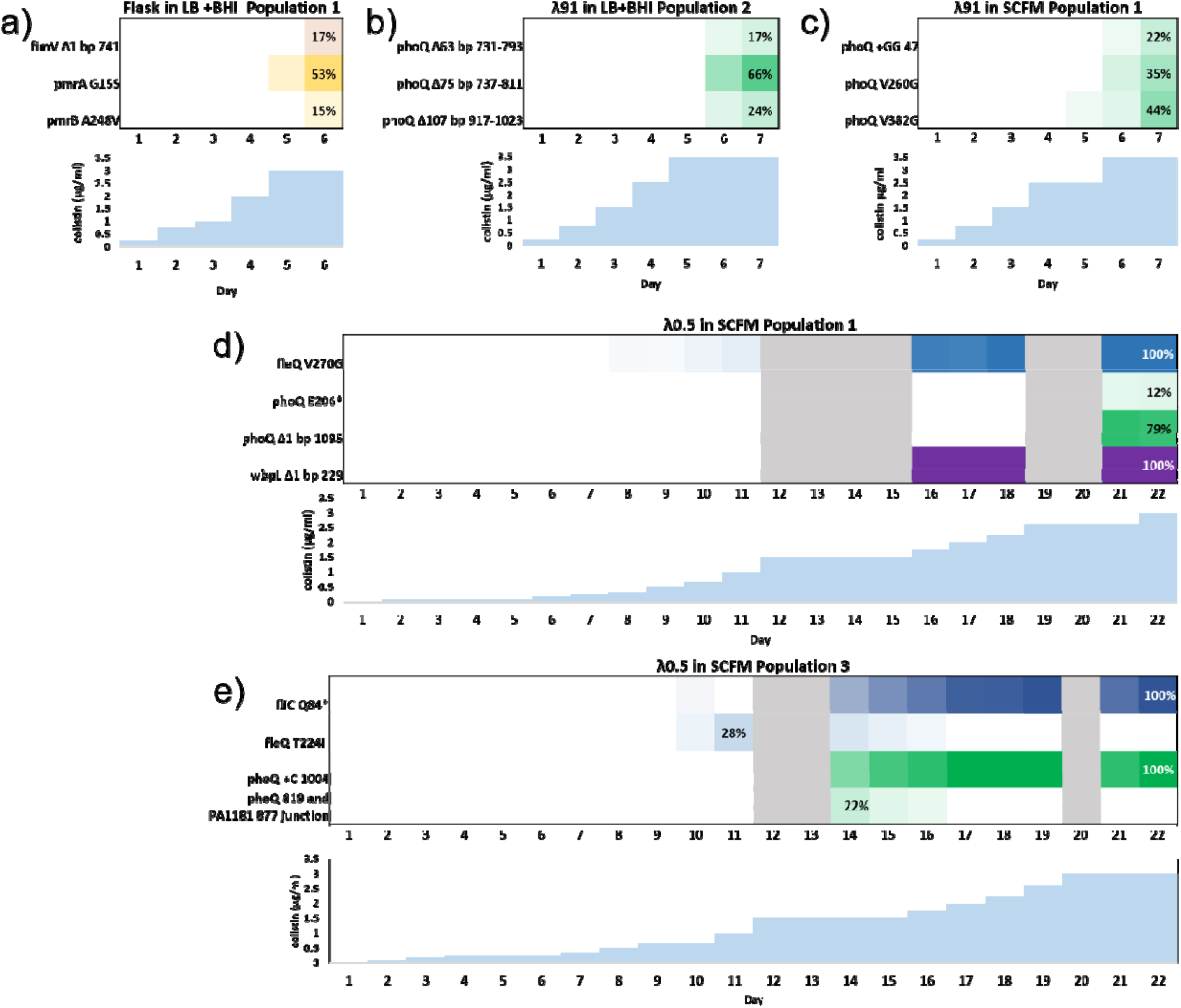
Longitudinal genomics for select populations from each λ condition show that adaptive allelic frequencies follow distinct evolutionary trajectories between conditions. Allelic frequencies on each day are represented by color scale, with colistin concentration in μg/m shown underneath. More intense color is used to represent a higher frequency of an allele in the population. Only mutations that were consistently present in the population above a 5% frequency are shown. Gray bars indicate days where too few cells were present for genomic sequencing due to the slow expansion of adaptive alleles in the λ0.5 populations. a) shows a representative flask population, b) shows a representative λ91 population evolved in LB+BHI, and c) shows a representative λ91 population evolved in SCFM. All three had quick adaptation times, taking around 6-7 days to grow in 3-3.5 μg/ml colistin. d) and e) show experimental populations from λ0.5 populations; both experienced multiple periods of low population density followed by expansion of new alleles to high frequencies.

### *pmrB* mutations emerge when populations isolated from microdroplets are reintroduced to a flask culture

To test whether adaptive mutants having success in the microenvironment of the microdroplets would also be successful in the batch conditions of the flask transfer environment, we grew subsets of mid to late stage microdroplet evolved populations in batch liquid culture conditions. On days when populations were experiencing slowed population expansion during the λ0.5 evolution, 1.5 ml of 0.0015 OD culture was decapsulated and grown overnight as stationary batch cultures at the same colistin concentration and media conditions.

When microdroplet populations were reintroduced to bulk culture, variants with *pmrB* mutations arose overnight. (**Fig. 5)**. That the same *pmrB* mutation emerged from resuspensions made on different days of the microdroplet evolution study suggested that the *pmrB* mutation was present for some time at undetectably low frequencies in the microdroplet population. When short read sequence data was inspected manually with Integrative Genomics Viewer (28), we confirmed that in λ0.5 Population 1, the *pmrB* mutation was present in the resuspensions on Days 14 and 15 at frequencies near 2% or lower from Day 16 onwards. The fact that it is present, but not being selected for despite conferring colistin resistance and that it readily increases in frequency in the resuspension, suggests that mutations in *pmrB* confer a greater fitness cost in microdroplet populations.

**Figure 4.**
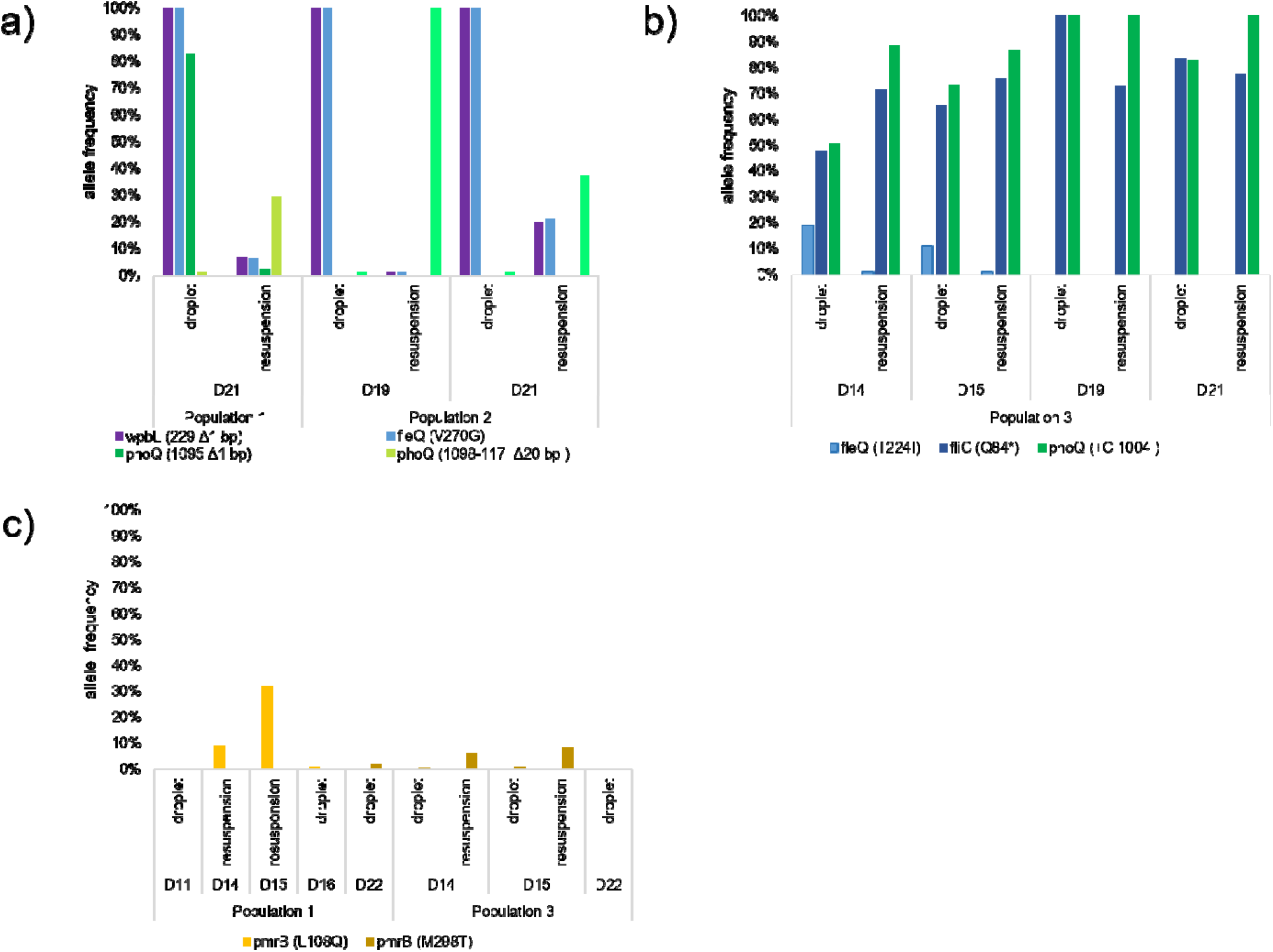
Alleles that were successful in λ0.5 microdroplet environments were less fit in well-mixed batch culture. λ0.5 populations on the indicated days were resuspended in a batch culture to assess their fitness. a) shows frequencies of adaptive *fleQ, wbpL*, and *phoQ* mutants in Populations 1 and 2; mutations in *fleQ* and *wbpL* decline dramatically in the resuspension cultures while a microdroplet *phoQ* mutation is replaced by an alternate allele in resuspension. b) shows allele frequencies of *fleQ, fliC*, and *phoQ* mutants in Population 3. Tracking allele frequencies over time of all three shows that an early *fleQ* mutation declines in both resuspensions and microdroplets, and is replaced by linked mutations in *fliC* and *phoQ*. After these mutations reached near fixation between Day 15 and 19, we observed that these two mutants maintained similar frequencies in microdroplets. In resuspensions the *fliC* mutation declined in frequency compared to the *phoQ* mutation. c) shows allele frequencies of *pmrB* mutants in Population 1 and 3. For some days the population was too small to estimate population size from OD and thus only resuspension or only microdroplet allele frequencies were available in some cases. Microdroplet data without corresponding frequency in resuspension is shown to illustrate baseline frequencies before and after resuspension data. While never observed *pmrB* mutations above 5% in microdroplets, we observed *pmrB* mutations emerge above 5% on multiple days in the resuspension.

**Figure 5.**
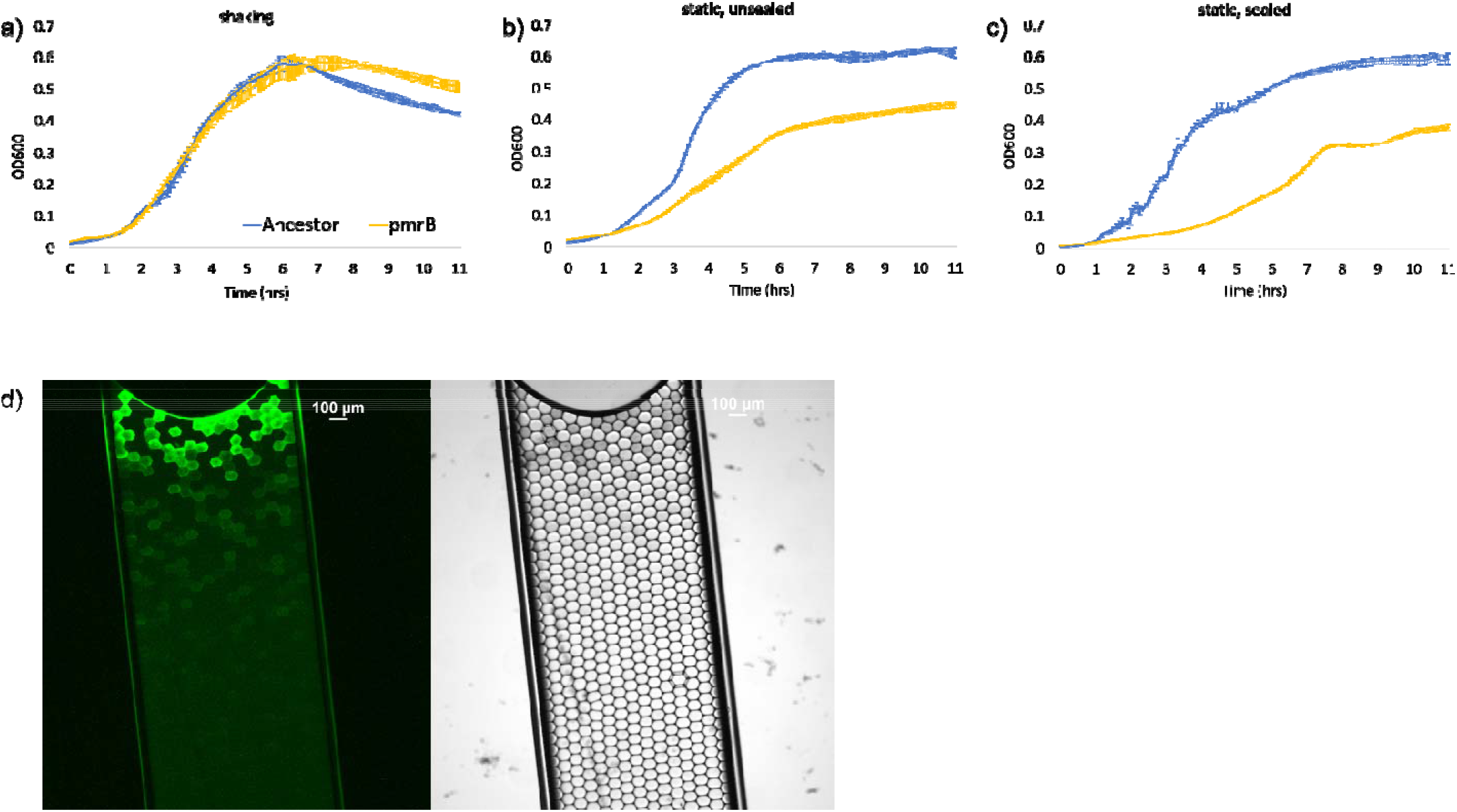
pmrB^(L108Q)^ grows slower than the ancestor in low oxygen conditions, which are present in microdroplets further from the air-droplet interface. a) PAO1 ancestor strain and pmrB^(L108Q)^ grow similarly in an aerated, well-shaken culture. b) pmrB^(L108Q)^ grows significantly slower than PAO1 in non-shaken culture, and c) shows that pmrB^(L108Q)^ grows slowest in a non-shaken culture with sides sealed with tape while PAO1 grows similarly in non-shaking, sealed, and in shaking conditions. d) images of GFP-labeled PAO1 grown in a capillary tube at 40X magnification are shown to illustrate the presence of an oxygen gradient across the aqueous layer of a microdroplet emulsion. The left image shows GFP fluorescence and the right image is brightfield microscopy. GFP activity, which relies on molecular oxygen, is high near the air-droplet interface but decreases as distance from the interface increases. The portion of the capillary tube is approximately .25 cm, 50% of the 0.5 cm average depth of microdroplet cultures during experimental evolution.

### *P. aeruginosa* PAO1 *pmrB*^*L108Q*^ has a reduced growth rate in the lower oxygen conditions that were present in the stationary microdroplet environment

To understand the lack of mutations in *pmrB* in microdroplet environments, we asked whether the distinct *pmrA* or *pmrB* evolutionary trajectories observed exclusively in flasks but not microdroplets reflected a particular environmental difference between the experimental systems. During experimental evolution, visual inspection of the aqueous phase of the microdroplet emulsion after overnight growth suggested that cells at the aqueous-air interface were growing faster than those at the oil layer. We reasoned that an important potential difference between the shaken batch flask culturing conditions and the stationary microdroplets might be the extent of oxygen diffusion. To test whether mutations to *pmrB* were at a disadvantage in microaerobic conditions, we introduced *pmrB*^*L108Q*^ into the PAO1 ancestor. *P. aeruginosa* PAO1 *pmrB*^*(L108Q)*^ was grown overnight in a 96-well plate under 3 different conditions; a well-shaken plate, a non-shaken plate, and a non-shaken plate with the sides sealed with tape to reduce oxygen exchange (**Fig 6**). While none of the conditions were anaerobic, the latter two plates approximated moderately aerobic and microaerobic conditions. The average growth rate of *pmrB*^*L108Q*^ during log phase in shaking culture was similar to the ancestor PAO1 strain (+0.15 OD/hr *pmrB*^*L108Q*^ vs +0.19OD/hr PAO1), but was ∼3-fold lower in static culture (+.080 OD/hr vs 0.26 OD/hr) and in static, tape-sealed culture (+0.05 OD/hr vs 0.17 OD/hr). The growth rate of *pmrB*^*L108Q*^ was significantly different between shaking and static, sealed conditions (p=0.003), while the growth rates of the ancestor PAO1 strain were not significantly different (p=0.14) (**S3**). Our data suggest adaptive mutations within *pmrB* during selection for colistin resistance confer slower growth in microaerobic environments, which may explain why *pmrB* mutations are not seen at high frequencies during the evolution of cells in microdroplets.

**Fig 6.**
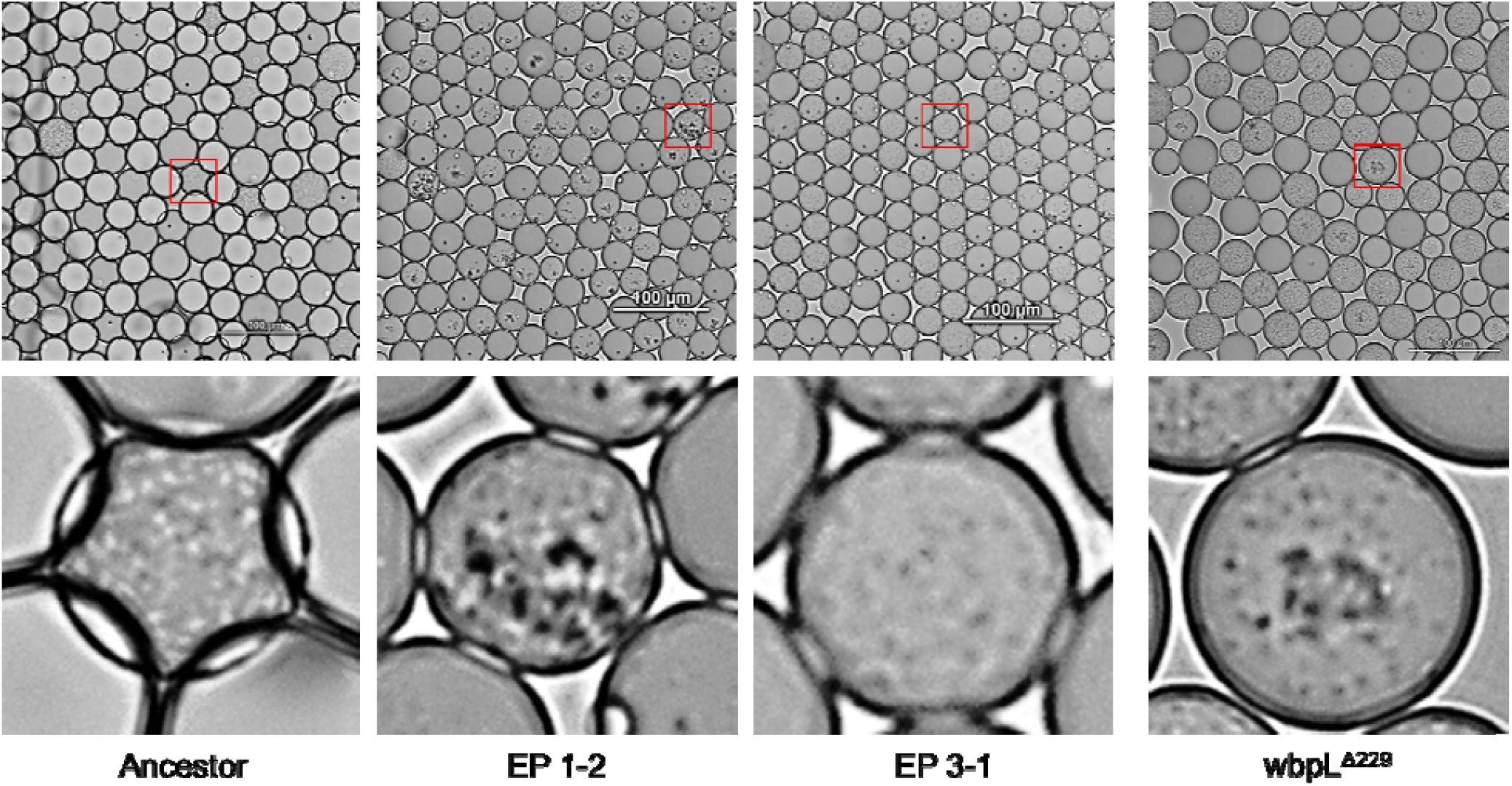
Isolates with adaptive mutations in *wbpL* show increased aggregation in pictures taken of different PAO1 isolates grown in microdroplet cultures for 24 hours under λ0.5 conditions. Bacteria appear as dark, small specks, while larger dark specks indicate bacterial aggregation. In the majority of populated microdroplets for the Ancestor and EP 3-1 (population contained *phoQ* and *fliC* mutations), bacteria are dispersed throughout the droplet indicating they are mainly planktonic. In the EP1-2 culture (population containing *wbpL, fleQ*, and *phoQ* mutations), the majority of microdroplets contain at least one aggregate, although not all bacteria appear to be in aggregates. In *wbpL*^(Δ229)^, several but not all microdroplets contain aggregates. This suggests that mutation in *wbpL* contributes to the aggregation phenotype seen in EP1-2, but other mutations in the population may increase the likelihood of aggregation.

### A loss-of-function mutation in *wbpL* affecting O antigen biosynthesis increased colistin resistance in microdroplet populations

From our longitudinal genomic data, we found mutations in *wbpL*, likely causing loss of O-antigen biosynthesis, were present only in λ0.5 populations evolved to colistin resistance. Two of the three experimental λ0.5 populations contained a single-nucleotide deletion at position 299(*wbpL*^(^Δ229)) at high frequency **(Fig 2a)**. The *wbpL* gene encodes for a glycosyltransferase that is required for synthesis of both types of O antigen present on lipopolysaccharides: the common polysaccharide antigen (CPA) (formerly A-band) and the O-specific antigen (OSA) (formerly B-band) (29, 30). The *wbpL*^(Δ229)^ in λ0.5 populations 1 and 2 is most likely a loss of function as it results in a frameshift mutation that deletes nearly three-fourths of the protein. A loss of *wbpL* function would result in the loss of both OSA and CPA on the outer membrane (30). Mutations in *wbpL*, and loss of O antigen biosynthesis more broadly is commonly observed in long-term CF isolates (5, 31). A recent study has also shown that O antigen may play a role in bacterial aggregate size and shape during infection as loss of O antigen increases cell surface hydrophobicity, leading cells to form dense clumped aggregates (29).

All three populations showed very slow expansion around 1.5 μg/ml; in Populations 1 and 2 *wbpL* mutations emerged at high frequencies during population re-expansion and phoQ mutations appeared near the end of the experiment at colistin concentrations comparable to when they were seen in λ91 populations (2.5 μg/ml) (**Fig 3e)**. In contrast, in Population 3 a *phoQ* mutation appeared directly after the number of viable cells declined at 1.5 μg/ml and a *wbpL* mutation did not appear. Taken together our data suggested that the wbpLΔ(229) mutation may have provided a low level of resistance to colistin, but likely to a lesser extent than a typical adaptive *phoQ* mutation. In order to confirm that *wbpL* loss of function was contributing directly to the colistin resistance observed in λ0.5 populations, we compared the colistin MIC of a *wbpL*^(Δ229)^ strain to that of the PAO1 ancestor and a *pmrB*^*(L108Q)*^ strain. *wbpL*^(Δ229)^ consistently grew at a two-fold higher colistin concentration than PAO1 (**Supplementary Figure 1, S4**).

### Multiple mutations affecting flagella function were observed in microdroplet populations

All λ0.5 microdroplet-evolved populations contained mutations affecting either flagella regulation or biosynthesis (**Fig 2)** through mutations in *fleQ* and *fliC*. Mutations in *fleQ* and *fliA* were observed in the λ91 SCFM control population, but not the experimental populations. In the λ0.5 control population there were additional mutations at a low frequency (between 5% and 20%) in *flgB, flgF, flgK, flhB, flhA*, and *fleS* (**S5**); *fleQ, fliC, fleS*, and the *flg* and *flh* genes are all involved in flagellar function. *fleQ* is involved in inversely regulating expression of flagellar biosynthesis genes and biofilm formation, and previous studies have found that SNP mutations in *fleQ* can result in aflagellate bacteria (32–36). *fliC* encodes the fliC flagellin protein that forms the main structural component of the flagella (37). The *fliC* mutation in λ0.5 population three resulted in a stop codon after less than a hundred nucleotides, very likely causing loss of functional flagella. *Flg* and *flh* genes encode for other proteins components needed for flagella function, including hook-associated flagellar protein (*flgK*) (38), flagellar basal body rod protein (*flgB, flgF*) (39), and transport into the periplasm (*flhB, flhA*) (40).

Flagellar mutants and subsequent changes in motility and exopolysaccharide production reproducibly evolve in laboratory biofilms and are also commonly seen in isolates from chronically infected CF patients (41–43). The appearance of these mutants at low colistin concentrations during evolution, as well as their appearance in the control, suggest they are an adaptation to the microdroplet environment rather than to the drug.

### Adaptive evolutionary trajectories identified in the late-stage of the microdroplet studies are not successful when reintroduced to flask culture conditions

In resuspension cultures, *fleQ* and *wbpL* mutant populations declined rapidly overnight, with the frequency in the population dropping from nearly 100% to 20-0% the next day (**Figure 4ab**). Over the same period of time, *fleQ* and *wbpL* mutations remained at high frequencies in the microdroplet populations carried out in parallel. In both Populations 1 and 2 of the resuspension conditions, *wbpL* and *fleQ* mutants declined simultaneously, making it difficult to disentangle the selection effects at play on each gene. However, in Population 3, the *fleQ* mutant frequency also decreased in resuspension compared to the microdroplet population in the absence of a *wbpL* mutation, indicating that the mutant *fleQ* is selected against on its own as well as in the presence of the *wbpL* mutant. The sharp decreases in frequency in resuspension cultures suggest that while *fleQ* mutants are advantageous in the microdroplet microenvironment, they confer a significant fitness cost in the flask environment.

In the resuspensions where *wbpL* and *fleQ* mutant populations are depleted, *phoQ* mutations that occurred in the *wbpL/fleQ* background are also depleted, but new phoQ mutations appear at high frequencies. Since the same *phoQ* mutation arises in subsequent resuspensions, this suggests that these *phoQ* mutants are present in the microdroplet populations at undetectably low frequencies, but are outcompeted by *fleQ/wbpL* or *fleQ/wbpL/phoQ* mutants (**Fig 4c)**. However, in the resuspension, *phoQ* mutants outcompete the *wbpL/fleQ* mutants. This dynamic further supports the idea that the *fleQ/wbpL* mutants have an advantage in the microdroplet microenvironment, but are at a disadvantage in the well-mixed flask cultures.

### Endpoint isolates with mutations within *fleQ* and *fliC* display a loss of motility; endpoint isolate from λ0.5 Population 2 and *wbpL*^(Δ229)^ display increased aggregation

To confirm whether the genotypes in the final populations conferred phenotypes associated with those mutations, we imaged endpoint isolates, *wbpL*^(Δ229)^, and *pmrB*^*(L108Q)*^ in microdroplets after 24 hours. Populations were imaged and recorded inside microdroplets after 24 hours of incubation to examine the spatial distribution and motility of the bacteria within the microdroplets. Images and recordings show that, as expected, the ancestor PAO1 strain is both planktonic and motile; bacteria within microdroplets are moving quickly and are evenly distributed throughout the microdroplets. Isolates 1-1 and 3-1 show loss of motility compared to the ancestor; movement within microdroplets is much slower (**S6**). Both of the populations from these isolates contained fixed flagellar mutations in either *fleQ* (Pop 1) or *fliC* (Pop 2), mutations that could cause loss of motility (35, 37). Isolate 1-1, from Population 1, containing a *wbpL*^(Δ229)^ and *fleQ*^*(V270G)*^ mutations at near fixation, and a *phoQ*^(^Δ1095) at 78%, shows the formation of clumped aggregates within microdroplets, which are not seen in either the ancestor or Isolate 3-1 which does not have the same mutations. *wbpL*^(Δ229)^ has a clumping phenotype that appears intermediate between Isolate 1-1 and the PAO1 ancestor, suggesting that *wbpL*^(Δ229)^ contributes to the clumping phenotype but interactions with the other mutations may exacerbate the phenotype.

## Discussion

*In vitro* experimental evolution provides an important approach to identifying genotypic changes and evolutionary trajectories responsible for increased antimicrobial resistance and there is increasing recognition of the importance of spatial structure on evolutionary outcomes (21, 44). Spatial structure can be provided by the environment itself, leading to multilevel partitioning of microbial communities (45) and by biofilm formation (7, 46). A conundrum confronted by investigators occurs when adaptive changes identified by *in vitro* studies provide either different evolutionary trajectories or a strong emphasis for the success of one trajectory over another that is not reflected in the clinical data. Logically, this is ascribed to the substantial differences between laboratory culture conditions versus the substantially more complex and nuanced environments of the patient or animal model. Experimental evolution using microdroplet emulsions provides an opportunity to more stringently control many aspects of the selection environment (22). In this study, we investigated the use of microdroplet-based experimental evolution to simulate the spatial structure that is present in most ecological and clinical contexts. In the CF patient lung, there are nearly infinite opportunities for localized (i.e., spatially structured) microenvironments that could provide different selection conditions, which can change over chronic infections that span years (47). Bacterial communities may be partitioned within airways and tissues that have varying oxygen microenvironments resulting from thick mucus and inflammation response (23, 48, 49).

In this study, we observed that evolving *P. aeruginosa* to colistin in a spatially segregated emulsion environment recapitulated both well-established resistance mutations and mutations commonly associated with long-term adaptation in a CF lung environment. Additionally, we observed that in contrast to batch suspension cultures where *pmrB* is the most common target for resistance mutations, in microdroplets, *phoQ* makes up th*e* most common adaptive allele in microdroplets.

### Selection against variants with changes in the two-component *pmrA/pmrB* two-component system in spatially structured microaerobic environments may replicate scenarios similar to that of the damaged lung tissues of CF patients

Mutations in *pmrB* and *phoQ* are often thought of as interchangeable in the context of colistin resistance, despite the number of differing transcriptional pathways each control (50). Clinical studies have shown that colistin resistance can arise from changes to either the *pmrA/pmrB* and *phoP/phoQ* two-component systems, even within the same patient (27, 51, 52). Here, we show that spatial segregation alters the selection pattern for *pmrB* mutants; *pmrB* mutations dominated in bulk flask cultures, but were present at very low levels in microdroplet populations and were only able to reach high frequencies when reintroduced to a bulk culture. We also showed that a *pmrB* mutant, *pmrB*^*(L108Q)*^ grew significantly slower compared to the ancestor PAO1 in microaerobic environments. When *pmrB*^*(L108Q)*^ was grown in well-shaken, non-shaken, and non-shaken sealed conditions, it exhibited decreased growth compared to the ancestor in the latter two conditions. Since microdroplet cultures are static and oxygen must diffuse from the top through multiple layers of the microdroplets (**Fig 5**), they are less well aerated than a well-mixed flask culture.

As mutations to *pmrA, pmrB*, and *phoQ* have been observed in clinical populations, the distinctiveness of the evolutionary trajectories identified in this study supports the notion that spatial structure within the lung leads to varying microenvironments favoring different evolutionary trajectories. Many microaerobic environments exist within the CF lung, and *P. aeruginosa* often undergoes significant transcriptional and genetic changes to adapt to these environments (53). A mutation in *pmrB* has previously been implicated in transcriptional dysregulation of *dnr*, a gene involved in regulating anaerobic nitrate metabolism (54), and the *pmrAB* TCS is involved in sensing and responding to extracellular Fe3+ which is common in aerobic environments (55, 56). *pmrB* is known to transcriptionally regulate a large number of genes, and it is possible that mutations in *pmrB* cause downstream transcriptional effects that dysregulate anaerobic metabolism and cause a slower growth of *pmrB* mutants in microaerobic environments. It is plausible that while both *pmrB* and *phoQ* mutants succeed in well aerated and moderately aerated regions of the CF lung, *phoQ* mutations dominate in microaerobic regions.

### Evolution in microdroplets reveals novel resistance pathways that recapitulate common adaptations to CF lung

In addition to changes within *phoQ*, multiple microdroplet populations had mutations in genes associated with loss of motility. Additionally, mutations in O antigen biosynthesis occurred in 2 out of the 3 experimental λ0.5 populations. Typically, loss of motility and loss of O antigen biosynthesis is thought to provide an adaptive advantage by increasing the bacterial cell’s ability to evade detection by the immune system (57). However, in our experiment, these mutations occurred in the absence of any type of immune system, and perhaps more significantly, the mutation in *wbpL*^(Δ229)^ was shown to confer a modest increase to resistance that was advantageous at the early stages of adaptation to colistin in microaerobic conditions.

Both loss of motility and O antigen biosynthesis have been implicated in biofilm formation and biofilm phenotypes and only arose in microdroplet environments (29, 41). Previous studies have shown loss of O antigen biosynthesis arises in laboratory biofilm evolution experiments, and that loss of O antigen biosynthesis can result in a denser aggregation phenotype that resembles aggregation phenotypes seen in CF patients (29). In addition, we observed that endpoint isolates taken from populations containing *wbpL* mutants also displayed increased self-aggregation in microdroplets. We speculate that loss of *wbpL* function and O antigen may increase self-aggregation, which then decreases bacterial exposure to colistin, which cannot efficiently penetrate the aggregates (29). These mutations, either individually or in concert, are clearly disadvantageous in well-mixed bulk culture but can be advantageous in noncompetitive, spatially segregated cultures. This supports that spatial structure is important for reducing competition and facilitating the growth of clinically relevant adaptation pathways.

### Microfluidic emulsions as a model for adaption to late-stage chronic infections

Overall experimental evolution to colistin resistance in microdroplet emulsions provided evidence that the success of particular evolutionary trajectories for adaptation to colistin through the two-component *pmrA/pmrB* or *phoP/phoQ* systems may be the result of spatial structure and oxygen levels within the CF lung. In addition, we showed that a mutation typically associated with chronic lung adaptation and immune evasion, wbpL loss of function, also confers colistin resistance. Our work highlights the importance of studying antibiotic resistance evolution in environments that may simulate relevant ecological conditions of the treatment environment and encourages the development of highly-controllable microfluidic emulsions to mimic the conditions during chronic CF lung infection or other niches of interest.

## Methods

We conducted multiple experimental evolutions in different media conditions with different levels of spatial segregation as shown in **Table 1**. Lambda is defined as the average number of cells/microdroplet (22). In total, we evolved three bulk flask populations in LB + BHI media, three populations with λ=91 in LB + BHI media, three populations with λ=91 in SCFM media, and three populations with λ=0.5 in SCFM media.

### Determining Spatial Segregation Parameters for Droplet Evolution

Serial dilutions followed by plate counts were used to calculate the conversion factor between OD and CFU for PAO1. The OD of the ancestor was measured after 24 hours of growth on a shaker at 37°C, then dilutions of 1×10^−6^ to 1×10^−10^ cells/ml were spread on LB agar plates. After 24 hours of incubation at 37°C, colonies were counted for plates that did not form a lawn. The calculated CFU averaged over four repetitions was 6.01×10^9^ cells/ml of culture at an OD=1.

### Droplet Generation Using Microfluidics Chips

Experimental evolution was conducted using two different levels of spatial segregation, which was controlled by changing droplet size and starting cell density. For λ=91 condition, cultures were diluted to OD=0.01 and encapsulated in 143 μm microdroplets using our microfluidic chip design (22). For the λ=0.5 condition, cultures were diluted to OD=0.001 and encapsulated in 46 μm microdroplets using the microfluidic chip design from Mazutis et al (58).

### Evolution in Microdroplets

SCFM media that mimics the nutritional composition of biological CF sputum was made and used for evolving the λ91 SCFM population and the λ0.5 populations (25). All populations were started from individual colonies of PAO1 with an MIC of 2 μg/ml colistin. 3 ml of media (SCFM or LB+BHI) were inoculated, encapsulated in microdroplets, and incubated in 50 ml conical tubes for 24 hours at 37°C without shaking. After 24 hours microdroplets were broken with a demulsifier (1H,1H,2H,2H-perfluorooctanol) and used to inoculate the next day’s culture with increased colistin. Colistin was increased over the course of evolution on the regimen shown in **Fig 3 and S2**; if growth was below 0.15 OD for the 143 μm microdroplets or below 0.075 OD for the 46 μm microdroplets, colistin concentration was kept at the previous day’s concentration until OD surpassed the threshold, at which point colistin concentration for the next day’s culture was increased again.

### Evolution in Flask

Cultures of PAO1 were made from colonies grown on LB plates and inoculated in LB+BHI broth. Cultures were grown for 24 hours at 37°C and shaking at 225 rpm. After 24 hours of growth, 1% (v/v) of the overnight culture was transferred to fresh growth media containing increased colistin.

### Collection of Endpoint Isolates

Once populations were able to grow at the desired final MIC, 100 μl from the final day’s population was streaked on LB agar plates and incubated at 37°C overnight. The next day, 3 colonies from each population were inoculated into SCFM media and grown overnight at 37°C and 225 rpm, then stored as glycerol stocks at -80°C.

### Measuring MIC of *wbpL*^(Δ229)^ and *pmrB*^*(L108Q)*^

To compare the colistin MIC of *wbpL*^(Δ229)^ with the PAO1 ancestor, we performed broth microdilution tests (testing growth in 0-16 μg/ml colistin). LB plates were streaked from glycerol stock and grown overnight. Colonies were then picked and grown overnight in SCFM. New cultures were inoculated and grown until cultures reached log phase. Cultures were diluted to 0.05 OD in new SCFM media on a 96 well plate and grown for 24 hours in a Spark Tecan at 37 °C.

### Growth on Congo Red Plates

To examine exopolysaccharide production and colony morphology, endpoint isolates were grown on VBMM Congo Red agar (41). Plates were incubated at 37 °C for 48 hours then imaged. Data is shown in **SI Figure 2**.

### Imaging Endpoint Isolates and GFP-PAO1 in Microdroplets

To examine the motility and aggregate formation of endpoint isolates in microdroplets, the ancestor and endpoint isolates 1-1 and 3-1 were incubated in 1.5 ml of 46 μm microdroplets in 50ml conical tubes at 37°C overnight without shaking. Microdroplets were transferred to capillary tubes (0.05 × 0.1 mm) and imaged at 200X magnification bright field microscopy with a Nikon camera.

To examine oxygen diffusion via GFP production, PAO1 with a chromosomal GFP gene was grown in 143 μm microdroplets in glass capillary tubes (0.05 × 0.1 mm) for 24 hours in triplicate in both 30°C and 37°C conditions. Capillary tubes were imaged at 40X magnification under both GFP fluorescent and brightfield microscopy with a Nikon camera.

### Whole Genome Sequencing and Analysis

After evolution was completed, DNA was extracted from the daily samples for all populations using Qiagen DNeasy Ultracean Microbial Kit following the manufacturer’s instructions. Illumina compatible libraries were prepared using plexWellTM 384 NGS multiplexed library preparation kit (seqWellTM), and pooled libraries for whole genome sequencing. Breseq software was used to assemble short read sequences and identify mutations (59). First, the short read sequences from Day 0 were aligned with a reference genome from NCBI (reference sequence NC_002516.2), and any differences were incorporated into a new reference using the gdtools APPLY function. Next this reference was used to align short read sequences from each of the daily populations; differences between the reference and the short read sequences were identified using the polymorphism flag -p with a 5% cut-off frequency for differences between the population sequences and the reference strain. Mutation tables across all days for a population were assembled using the gdtools COMPARE function. Any mutation that appeared at greater than 15% frequency on two or more days was considered significant and included in the final data. If significant mutations had gaps between days where mutations were present, the short read sequences were examined in Integrative Genomics Viewer (IGV) and mutation frequency was calculated manually (28). Raw sequence data is available on the NCBI Sequence Read Archive (SRA) database (BioProject ID PRJNA978079).

### Constructing wbpL and pmrB single mutants

Single mutants in PAO1 were constructed using the protocol described by Mehta *et al* (13). The *wbpL* and *pmrB* single mutants were generated from the ancestor PAO1 strain. Electroporation of the constructed plasmid into PAO1 and the strain selection was performed as described previously (13). The mutants were cultured, and the DNA was sequenced using Sanger sequencing to identify the mutations.

### Testing Growth of *pmrB*^*(L108Q)*^

Colonies of *pmrB*^*(L108Q)*^ were picked from LB plates and grown overnight in SCFM. After performing 1% dilutions and growing until new cultures reached log phase, new cultures were made with a concentration of 0.01 OD per 0.1ml on a 96 well microplate and grown for at least 15 hours on a Tecan (Spark) microplate reader at 37°C or Epoch 2 microplate reader. Cultures were either grown shaking (linearly 450 rpm), non-shaking/static, or non-shaking with taped sides.

## Author Contributions

Y.S., H.M., and S.D. conceived of the project. S.D., H.M. S.S., and Y.S. designed the experiments. S.D., S.S., and K.G. performed all the experiments. S.D. and Y.S. wrote the original manuscript. All authors reviewed and commented on the manuscript.

## Acknowledgments

The authors acknowledge the use of resources of the Shared Equipment Authority at Rice University for this work. We are thankful to Cailey Renken for assistance during conducting the flask transfer experimental evolution of PAO1 to colistin. This work was supported by NIAID Award R01A1080714 to Y.S. The figures and schematics were created with MATLAB, GraphPad Prism 9, BioRender.com, Excel, and Powerpoint.

