## Supplementary material for "Experimental evolution of *Pseudomonas aeruginosa* to colistin in spatially confined microdroplets identifies evolutionary trajectories consistent with adaptation in microaerobic lung environments": SI

1. Longitudinal mutation data for all populations is compiled in significant_mutations_for_all_populations.xlsx (**S5**)
2. Data on OD and colistin concentration during experimental evolution for all populations is compiled in allpops_experimental_evolution_colistin concentration_and_OD_by_day.xlsx (**S1**)
3. Growth curve data and growth rate calculations for the PAO1 Ancestor and pmrB(L108Q) in shaking, non-shaking, and non-shaking sealed conditions are compiled in pmrB_and_ancestor_growth_curves.xlsx (**S3**)
4. MIC broth microdilution data is available in the excel file MIC_test_PAO1_wbpL_&pmrB.xlsx **(S4)**
5. Videos showing the motility within droplets of the PAO1 Ancestor, EP1-2, EP1-3, wbpL, pmrB are accessible in the folder microdroplet_videos. (**S6**)
6. Toy model of lambda expansion (**S2)**

**Supplementary Figure 1** Broth microdilution test of growth of PAO1, *wbpL*$^{(\Delta 229)}$, and *pmrB*$^{(L108Q)}$ at 0, 0.5, 1, 2, 4, 6, 8, and 16 µg/ml colistin shows that mutant *wbpL* grows at twofold higher colistin concentration than the ancestor

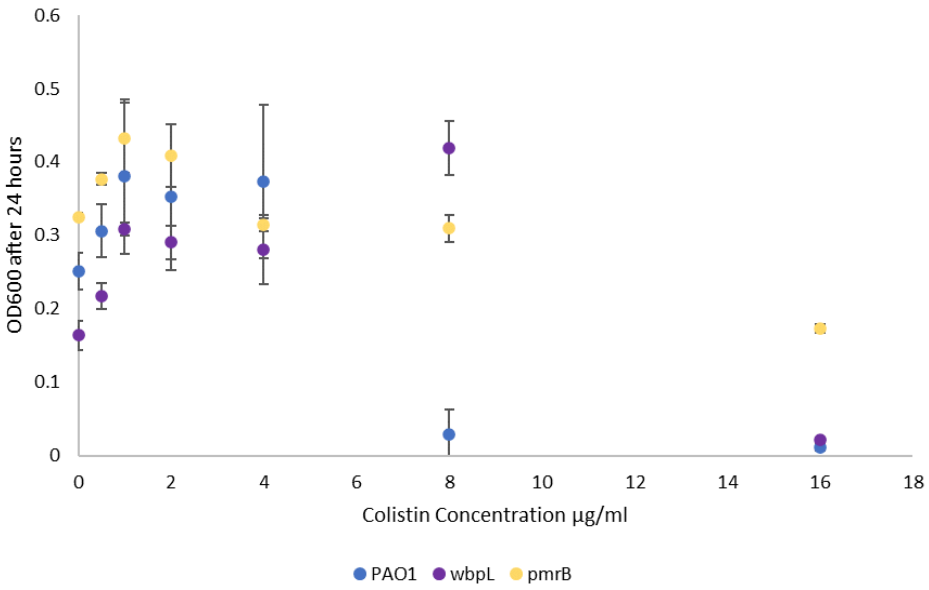


PAO1 strain. Graph shows averaged final OD after 24 hours of continuous growth for each strain at varying colistin concentrations.

**Supplementary Figure 2** shows differing colony morphologies of the PAO1 Ancestor, EP 1-2 (population containing high-frequency mutations *phoQ*, *fleQ*, and *wbpL* mutations), and EP3-1 (population containing high

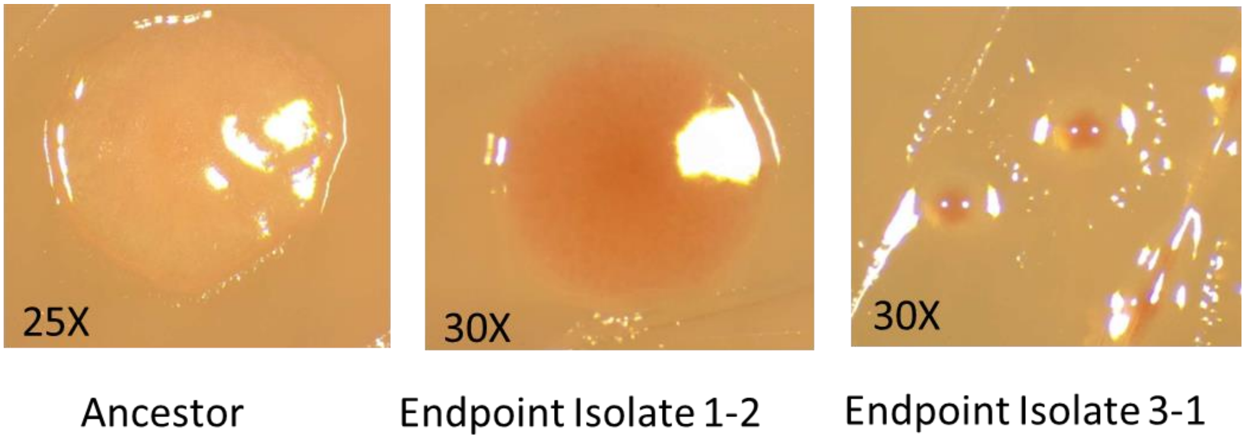


frequency *phoQ* and *fliC* mutations). Typically, increased absorption of Congo Red dye indicates increased exopolysaccharide production.
